# Mobile DNA Activity in Parkinson’s Disease: A Locus-Specific View of Endogenous Retroviruses

**DOI:** 10.64898/2026.07.03.736370

**Authors:** Elizabeth Banda-Arnold, Charles S. Venuto, Keith A. Crandall

## Abstract

Human endogenous retroviruses (HERVs) are mobile genetic sequences derived from ancient retroviral infections. While typically silenced, their reactivation has been implicated in gene dysregulation, aging, and immune-related transcriptional pathogenesis of some neurodegenerative diseases. Parkinson’s disease (PD) is the second most common neurodegenerative disorder, yet its etiology and HERV reactivation remain poorly understood. This study investigates locus-specific HERV expression in early-stage PD, including genetic and non-genetic cases (all PD), idiopathic PD without a known genetic cause (iPD), and PD driven by leucine-rich repeat kinase 2 mutations (LRRK2 PD). We analyzed RNA-seq whole-blood samples from 492 individuals (358 all PD, 256 were iPD, 63 LRRK2 PD, and 134 healthy controls (HC)). We identified 20 significantly dysregulated HERV loci in all PD versus HC. Five HERV loci were shared with iPD analysis, and one HERV locus was shared with LRRK2 PD. Notably, these shared loci included HERV-H and ERVLE elements, indicating robust disease-associated retroviral signals independent of disease subtype. We found that genes proximal to these HERVs revealed pathways implicated in PD pathogenesis. Immune cell deconvolution showed increased neutrophil abundance and decreased resting CD4+ memory T cells proportions across the PD cohorts when compared to HC, consistent with neutrophil-lymphocyte ratio observed in previous peripheral immunity studies. Transcriptomic HERV alterations are present in whole blood across PD populations and are associated with dysregulation of fundamental cellular pathways and peripheral immune remodeling. Our findings motivate experimental validation of locus-specific HERV expression as a candidate blood-based signature with potential to inform PD neuroinflammatory and neurodegenerative processes.

## Introduction

Human endogenous retroviruses (HERVs) are a subgroup of transposable elements (TEs) that comprise ∼8% of the human genome^1^. These fossil retroviral remnants in human DNA sequences are the cumulative result of viral infections that became fixed and integrated in the host genome during evolution. HERVs belong to the subtype of Long Terminal Repeat (LTR) retrotransposons, meaning they spread by a ‘copy-paste mechanism’ also known as replicative transposition^2,3^. HERVs share structural features with exogenous retroviruses organized as 5′ LTR – gag – pol – env – 3′ LTR, but most copies are highly mutated, truncated, or present only as solo LTRs^4^. The reactivation process is particularly associated with younger HERV subfamilies, such as HERV-K, HERV-W, and HERV-H^5^. These younger elements retain a more intact open reading frame regardless of accumulated mutations, enabling them to reactivate and produce functional viral components.

Although the majority lack protein-coding potential, they contain functional cis-regulatory elements, including promoters, enhancers, and polyadenylation signals, which allow them to modulate the transcription and functions of neighboring host genes. Additionally, ERV loci, especially solo LTRs, can serve as transcriptional start sites for long non-coding RNAs (lncRNAs). Studies exploring aging and chronic inflammation have implicated the reactivation of HERVs as a significant contributor to pathogenic burden in chronic illnesses, viral infections, and neurodegenerative diseases^6–8^. The mobilization may result in mutagenesis that disrupts the open reading frame of coding or regulatory regions. In healthy tissues, expression is tightly modulated and silenced by epigenetic mechanisms and cellular homeostasis, including DNA methylation and histone modifications. Derepression can occur in response to genetic variation, environmental triggers, cellular stress, and aging-associated epigenetic erosion. Aberrant ERV reactivation has been associated with DNA damage, modulation of immune genes, and cytotoxic effects^2,7^. A well-described example of ERV-mediated immune interaction is the regulation of interferon-gamma–responsive genes (IFN-γ), such as AIM2, APOL1, and IFI6, by MER41B elements. MER41B elements are located near the AIM2 gene and, upon IFN-γ stimulation, drive transcriptional activation of adjacent immune genes, providing beneficial signals and innate immune system response^9,10^. However, chronic or dysregulated activation of these elements contributes to pathological immune states and autoinflammatory conditions.

In recent years, this knowledge has driven interest in the development and progression of neurodegenerative diseases such as multiple sclerosis (MS)^8^, amyotrophic lateral sclerosis (ALS)^11^, and Alzheimer’s disease (AD)^12^. Among these, Parkinson’s disease (PD) is the second most common neurodegenerative disorder globally, with its prevalence and disability rates increasing more rapidly than AD^13^. The global burden of PD is expected to surge by 112% between 2021 and 2050, with case numbers projected to reach 25.2 million. PD is a complex multifactorial condition whose cause is associated with both environmental and genetic risk factors. Currently, its precise pathogenesis is not fully known, but it is characterized by the aggregation of misfolded alpha-synuclein, resulting in the formation of Lewy bodies in the brain and dopaminergic neuron death^14^. Additionally, peripheral inflammation, molecular changes, and systemic transcriptional analysis patterns have revealed correlations with clinical features, ongoing microglial activation, and accelerated disease progression^15–17^.

Research on the potential role and importance of ERV proviral loci in PD etiology is sparse and has thus far solely focused on family-level analysis such as HERV-K or PD as a mixed cohort. One HERV study found HERV-K was significantly downregulated in the postmortem pre-frontal cortex and blood in late-stage PD patients, with its presence in astrocytes^18^. Additionally, in both the brain and blood, HERV-K was associated with lower levels of GFAP, a marker of astrocyte health and damage. These findings correlated with disease severity and progression, suggesting a loss of a potentially protective astrocyte function.

Overall, studies have shown HERV-K expression correlates with genes involved in key PD pathways, including apoptosis, cellular senescence, mitochondrial dysfunction, and alpha-synuclein related pathology^2,18–20^. The HERV-K envelope (env) protein has been implicated in astrogliosis, and viral particles may promote neuroinflammation in the presence of alpha-synuclein fibrils. Both of these processes contribute to dopaminergic neuron vulnerability and disease progression. This highlights the potential of HERV derepression and warrants further investigation on a locus level. Locus-specific investigation will address the limitations of HERV family heterogeneity and provide insights to consider the individual insertions and structures of these unique markers.

Since transcriptional changes in blood can provide insightful clues for PD pathogenesis, in this study, we analyzed RNA-seq whole-blood samples from 492 individuals with early-stage PD. These include genetic and non-genetic cases (all PD), idiopathic PD without a known genetic cause (iPD), and PD associated with leucine-rich repeat kinase 2 mutations (LRRK2 PD), and healthy controls (HC). We utilize Telescope^21^, a computational software tool that leverages a Bayesian mixture model^22^ combined with an expectation-maximization algorithm to assign mapped RNA-sequencing (RNA-seq) fragments to their locus of origin. We provide insights into the retrotranscriptomic profile of HERV expression in PD, shedding light on the molecular mechanisms and peripheral inflammation in PD.

## Methods

### Participant characteristics

We obtained clinical information of RNA-seq data from the Michael J. Fox Foundation Parkinson’s Progression Markers Initiative Data Repository. Data used in preparation of this article was obtained on [26-06-13] from the Parkinson’s Progression Markers Initiative (PPMI) database (www.ppmi-info.org/access-data-specimens/download-data), RRID:SCR_006431. For up-to-date information on the study, visit www.ppmi-info.org. Our study has a total of 492 participants (358 cases (all PD), and 134 HC) reported in Supplementary Table 1. PD participants consisted of a higher proportion of males (208; 58.2%) compared to females (150, 42%). The average baseline age of participants was 61.7 years (S.D. 9.8). In terms of racial demographics, the majority were White (337, 94.13%), followed by Black (5, 1.4%), Asian (4, 1.1%), Other/Multiracial (11, 3.1%) and 1 unknown. Based on years since diagnosis and Hoehn & Yahr Stage Score most participants were primarily in the early stage of disease: Score 1 (140,40%), Score 2 (207,58%), and a small number with Score 3 (7, 2%). The HC participants held a similar ratio of 96 males (64%) and 48 females (36%). The average baseline age was 60.22 years (S.D. 11.42). The majority were also White (123, 92%), followed by Black (7, 5.2%), Asian (1, 0.7%), other/Multiracial (3, 2.2%). All HC held a score of 0 on the Hoehn & Yahr Stage Scale indicating no signs of disease. For the additional analysis to access overlapping HERV loci, participants were also stratified by subgroups based on the presence or absence of manifest PD risk genes: *LRRK2* (n=84), *LRRK2* + *GBA* (n=7) and Sporadic (n=256).

### Clinical and RNA-seq data

Paired-end RNA-seq data (150 bp reads from an Illumina NovaSeq6000 platform) was obtained from The PPMI through Aspera downloaded in FASTQ and BAM format as part of the RNA Sequencing Project ^23^. In the PPMI whole blood RNA-seq project, samples were sequenced to an average of ∼100 million paired-end reads to create a high-quality transcriptome dataset. The files contained transcript abundance estimates presented as both featureCounts^24^ and Transcripts per Million (TPM) calculated using the Salmon method^25^. RNA-seq metadata for PPMI participants was provided in the file metaData.tsv, containing participant identifiers (PATNO), disease status, clinical event, and sequencing waves information. We selected 492 subjects at baseline from Phase 1 patients in the following diagnosis: Genetic PD, Idiopathic PD, Healthy Control and Genetic Unaffected. Comprehensive clinical data was obtained from the PPMI Clinical Study Curated Data Cut_20251112. To ensure accurate participant classification of ‘condition’ factor during downstream analysis, individuals were categorized as PD cases or HC based on PPMI predefined COHORT and subgroup variables in PPMI Clinical Study Curated Data Cut_20251112. iPD was used for “Idiopathic PD” or “Sporadic PD”, LRRK2 PD was used for “*LRRK2*” and “*LRRK2* + *GBA*” then merged as one genetic group for analysis labeled “LRRK2 carrier”. For HC versus LRRK2 carriers, analysis was limited to plate 1-27 to account for disproportionate case-to-HC ratio on plate 28. This excluded 3 cases and 27 LRRK2 PD from the analysis.

### Quality control

To prepare the FASTQ files, we merged data across the different lanes into a single sample R1 and R2 file employing 4 parallel threads. We then performed FastQC to assess quality and contamination on MultiQC. Filtering and selection was performed using UNIX command-line tools to generate a subject list that matched both metadata criteria (plate 1-28, pass qc FLAG) through ∼4000 BAM files. This criteria excluded samples with quality control (QC) issues such as gender mismatches, PCA, outliers and low read counts in Phase 1 along our selected groups^23^. We obtained QC metrics and alignment mapping statistics of BAM files using samtools flagstat function and Python3. Trimming was not performed, as BAM files and count files were generated using STAR v2.6.1, a modern aligner which employs soft-clipping on low-quality bases and Salmon v0.11.3, a pseudo-aligner tool that does not require trimming for accurate quantification. This approach preserves read coverage and minimizes data loss.

### Retrotranscriptome quantification

BAM files were sorted by read name prior to retrotransposon quantification to ensure compatibility with downstream analysis. Locus-specific HERV expression was then estimated using Telescope^21^. Telescope deals with the ambiguous mapping of repetitive TE sequences by employing the Bayesian mixture model and expectation-maximization algorithm to reassign ambiguously mapped RNA-Seq fragments to the likely locus of origin, thereby facilitating accurate, locus-specific HERV quantification^22^ . Reads were reassigned using the default parameters (max_iter = 100, theta_prior = 200000). There are three types of output generated by Telescope, transcript counts derived from the EM-based reassignment, a statistical report summarizing each run, and an optional updated SAM file reflecting read reassignments.The count file represents inferred TE activity based on the detection of hallmark genomic features, including 5′ and 3′ LTRs flanking an internal open reading frame, which together indicate potentially functional elements. These counts were used for downstream analyses.

### Host gene expression analysis

To characterize transcriptional differences between all PD versus HC, we conducted differential expression analysis on RNA-seq gene-level counts formatted into a numerical matrix and the sample metadata. Gene identifiers were mapped to chromosomal locations using biomaRt (Ensembl, *Homo sapiens* dataset^26^). Genes with FDR adjusted *P* < 0.05 and shrunk |log_2_FC| > 0.1 were considered significantly differentially expressed. The apeglm algorithm^27^ was used similarly to the HERV differential expression. Principal Component Analysis (PCA) was performed using the plotPCA function in DESeq2.

### Cell type abundance

CIBERSORTx^28^ was used to isolate and characterize cell-type abundance from gene expression profiles in bulk RNA. Output cell-fraction estimates are used for downstream analyses to characterize immune composition and evaluate specific immune subsets contributing to observed transcriptional differences between the various PD groups compared to HC. We generated CIBERSORTx-compatible input files from DESeq2 normalized gene-level counts. Normalized counts were merged with the HUGO Gene Nomenclature Committee (HGNC) symbol and reformatted to the CIBERSORTx “mixture” format. Because multiple Ensembl IDs can map to the same HGNC symbol, counts were aggregated to the gene symbol by summing expression values across all Ensembl entries sharing the same HGNC symbol. Deconvolution was performed using the LM22 leukocyte signature matrix with 1,000 permutations, verbose output enabled, and quantile normalization disabled (QN = FALSE), as recommended for RNA-seq data. To focus on the most biologically relevant and reliably detected populations, the CIBERSORTx-derived immune cell fractions were filtered to retain samples with deconvolution *P* < 0.05 and with major innate and innate and adaptive immune populations previously implicated in systemic immune alterations, including neutrophils, monocytes, CD4 T-cell subsets, NK cells, and naive B cells. Beta regression models were fit for the seven most dominant immune cell types to test differences between PD groups and HC while adjusting for condition, age at visit, sex, and BMI. *P* values were corrected using the Benjamini–Hochberg method, then significant cell types (FDR-adjusted *P* < 0.05) were further analyzed.

### Differential expression analysis

To reduce sex-specific variance, features annotated as Y-chromosome-linked based on transcript pattern “Yq” or “Yp” were removed (n = ∼700) from the counts data prior to downstream analysis. Differential expression analysis was conducted comparing all PD to HC, iPD to HC, and LRRK2 PD to HC blood samples using DESeq2 v1.50^29^. The count files generated by Telescope were imported into R and compiled into a single matrix and then loaded into DESeq2 using the DESeqDataSetFromMatrix function. Age at visit, sex, BMI and condition (i.e., PD status) were incorporated into the DESeq model. A missing BMI value (n = 1) was imputed using linear regression based on age and sex of available samples. Continuous variables were standardized, and categorical variables were modeled as factors with controls as the reference level. The analysis applied a negative binomial model to account for the overdispersion of count data, identifying differentially expressed transposable elements with a significance threshold of adjusted *P* < 0.05. To obtain stable and interpretable effect size estimates, log_2_ fold changes were shrunk using the apeglm algorithm^27^ implemented in lfcshrink. Shrunk log_2_ fold changes were used for effect size ranking and visualization, while statistical significance was determined from the unshrunk DESeq2 results. For downstream reporting of biologically meaningful changes, loci were classified as differentially expressed if they satisfied both false discovery rate (FDR) adjusted *P* < 0.05 and shrunk |log_2_FC| > 0. Results were visualized with ggplot2 v 4.0^30^ using volcano plots to highlight elements with significant expression between all PD and HC participants, and a histogram to visualize the genomic distribution of differentially expressed loci. Following cell-type deconvolution, estimated proportions of neutrophils and resting CD4+ memory T cells were incorporated as covariates in differential expression analyses to evaluate the influence of cell-type composition on HERV expression.

### Proximal gene set enrichment and pathway analysis

HERVs have also been proposed as regulators that act on proximally-located genes^31,32^. We performed gene set enrichment analysis of those genes which were in the genomic neighborhood of the differentially expressed HERVs that were revealed in our analysis. Using an R script and Telescope HERV annotation (https://github.com/mlbendall/telescope_annotation_db), differentially expressed HERVs were mapped to their closest upstream, downstream and in some cases intersecting genes. A ranked gene list was generated using the shrunk logfold change × FDR adjusted *P* to prioritize HERV loci both statistical significance and HERV differential expression. Multiple HERVs mapping to the same gene were collapsed to retain the largest score. The ranked gene list was analyzed using the gseGO function in the clusterProfiler v4.18.8 R package^33,34^, which employs a permutation-based approach to estimate the null distribution of enrichment scores. Gene set size thresholds were set to a minimum of 10 and a maximum of 800 HERV loci per family. To ensure reproducibility, a random seed was set prior to analysis.

Running enrichment scores were calculated without numerical adjustment (eps = 0), and multiple hypothesis testing was corrected using the Benjamini–Hochberg method FDR to preserve the ranking of gene scores. Enrichment and significance of results were evaluated using normalized enrichment score (NES), core enrichment and adjusted *P*.

## Results

### Dataset

Across all samples, the median number of primary reads was ∼231M and minimum was ∼180M. The median mapping rate and proper pairing was 92.6% indicating high-quality alignments for downstream analysis.

### HERV expression

Telescope quantified 26,362 distinct loci expressed across the all PD versus HC cohort in our dataset and identified 45 retrotransposons loci with 20 HERVs and 25 Long Interspersed Element-1 (L1) as significantly differentially expressed (FDR adjusted p-value <0.05) (see Fig. 1 for HERV DE; and see Supplementary Table 2 for all retrosponsons). Among the HERVs, 13 upregulated and 7 downregulated with HERV-H elements dominating the upregulated side. In the iPD versus HC analysis, we identified 15 retrotransposon loci, 8 L1 elements and 7 HERV elements (6 upregulated and 1 downregulated) as shown in Supplementary Table 3. In LRRK2 PD versus HC, we identified 3 retrotransposon loci, 2 L1 elements, and 1 HERV (upregulated) as shown in Supplementary Table 4. When assessing overlapping HERVs across various expression, in all PD and iPD, we identified 5 overlapping loci, 1 downregulated loci ERVLE_8q12.3f and 2 upregulated HERVH_4q31.1a and HERVH_5q23.2g, 1 HERVL_10q11.23, 1 HERVFH19_2p22.2. 2 upregulated loci did not overlap, ERV316A3_12q24.13 and HERVH48_7q22.1 (Supplementary Fig. 1a). In all PD and LRRK2 PD, we identified 1 overlapping upregulated HERV locus ERVLE_3q25.32 (Supplementary Fig. 1b). To assess the impact of cell-type composition, we included estimated proportions of neutrophils and resting CD4+ memory T cells as covariates. This adjustment eliminated differential expression of HERV loci; however, in a reduced model including cell-type proportions and condition only, one locus (ERVLE_8q12.3f) remained significant, consistent with our primary analysis.

**Fig. 1:**
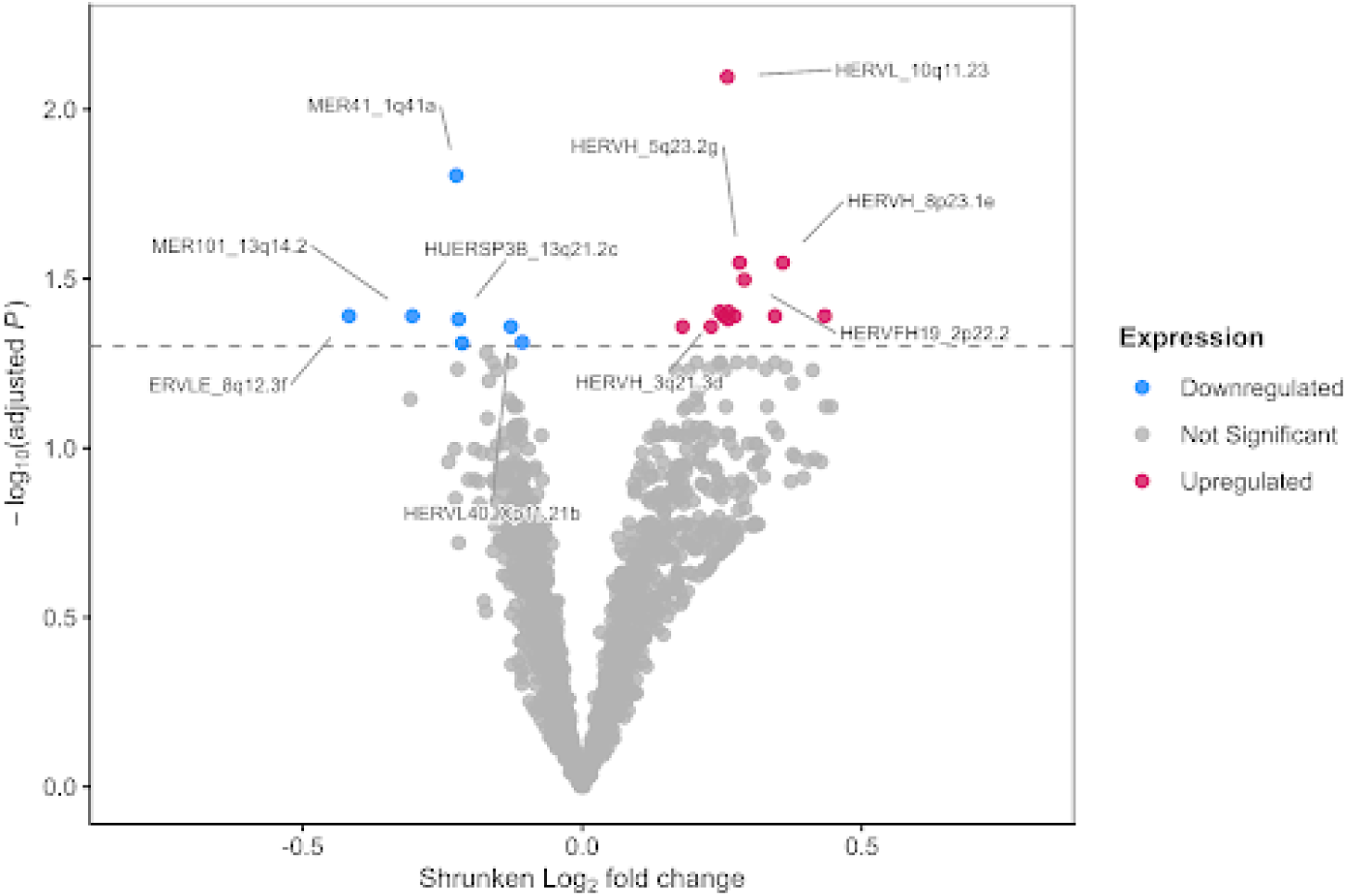
Differential expression of HERV Loci in all PD versus HC. Volcano plot of differential expression HERV analysis of all PD compared to HC. Points in the volcano plot are individual loci, their X-axis position represents the log_2_ fold change after apeglm shrinkage and the Y-axis position represents the –log_10_ of the FDR-adjusted *P.* The labeled dots represent HERV loci with the five most significant features per expression group (upregulated and downregulated), ranked by significance and effect size.

To further investigate the genomic distribution of differentially expressed HERV loci, we generated a barplot mapping their chromosomal locations. The plot illustrates loci across the genome, with upregulated loci and downregulated loci localized across multiple chromosomes.

The 20 HERV loci in all PD and vs HC were distributed across 14 chromosomes and a predominance of q arm localization (65%). HERV-H was the most represented, with 7 loci all upregulated across chromosomes 3q, 4q, 5q, 8p, 11q, 12p, and 14q. ERVLE showed mixed regulation, with one locus upregulated (3q25) and two downregulated (8q12, 1p32). All three MER elements (MER41, MER101, MER4B) were downregulated and distributed across the q arm of chromosomes 1,12 and 13. Notably, MER101 and HUERSP3B both map to chromosome 13q, making it the only chromosome with two downregulated elements. In iPD versus controls, 7 HERV loci were across 5 chromosomes. HERV-H again dominated with 3 upregulated loci (4q31, 5q23, 7q22). In LRRK2 carriers versus controls, ERVLE_3q25.32 was located on chromosome 3 (Fig. 2a, b).

**Fig. 2:**
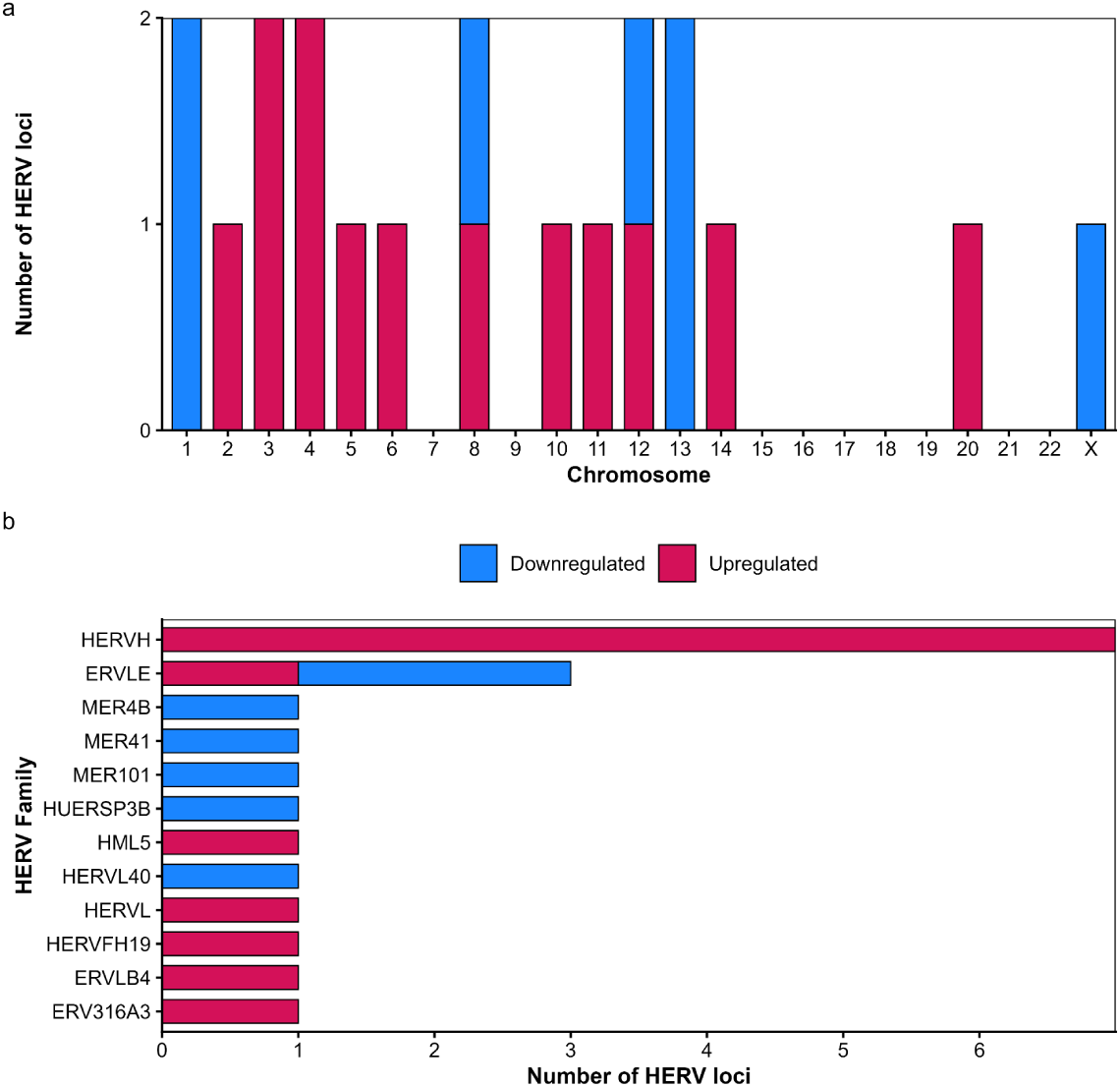
Differentially expressed HERV loci chromosome and family-specific patterns. **(a)** Bar chart showing the number of differentially expressed human endogenous retrovirus (HERV) loci across chromosomes. **(b)** Horizontal bar chart depicting the distribution of differentially expressed HERV loci by HERV family. In both panels, blue bars indicate downregulated loci and pink bars indicate upregulated loci.

HERVs can be divided into evolutionary families that exhibit unique expression patterns^35^. We sought to identify which families were disproportionately represented among the differentially expressed loci by calculating the HERV ratio within each family relative to its total detected loci. Although HERVs were distributed across 12 families, our analysis revealed that ERVLE and ERV316A3 families had the highest overall representation.(Fig. 3b). Few families show notable differential signals, HERV-H exhibiting the strongest overrepresentation in all PD (Fig. 3a-c).

**Fig. 3:**
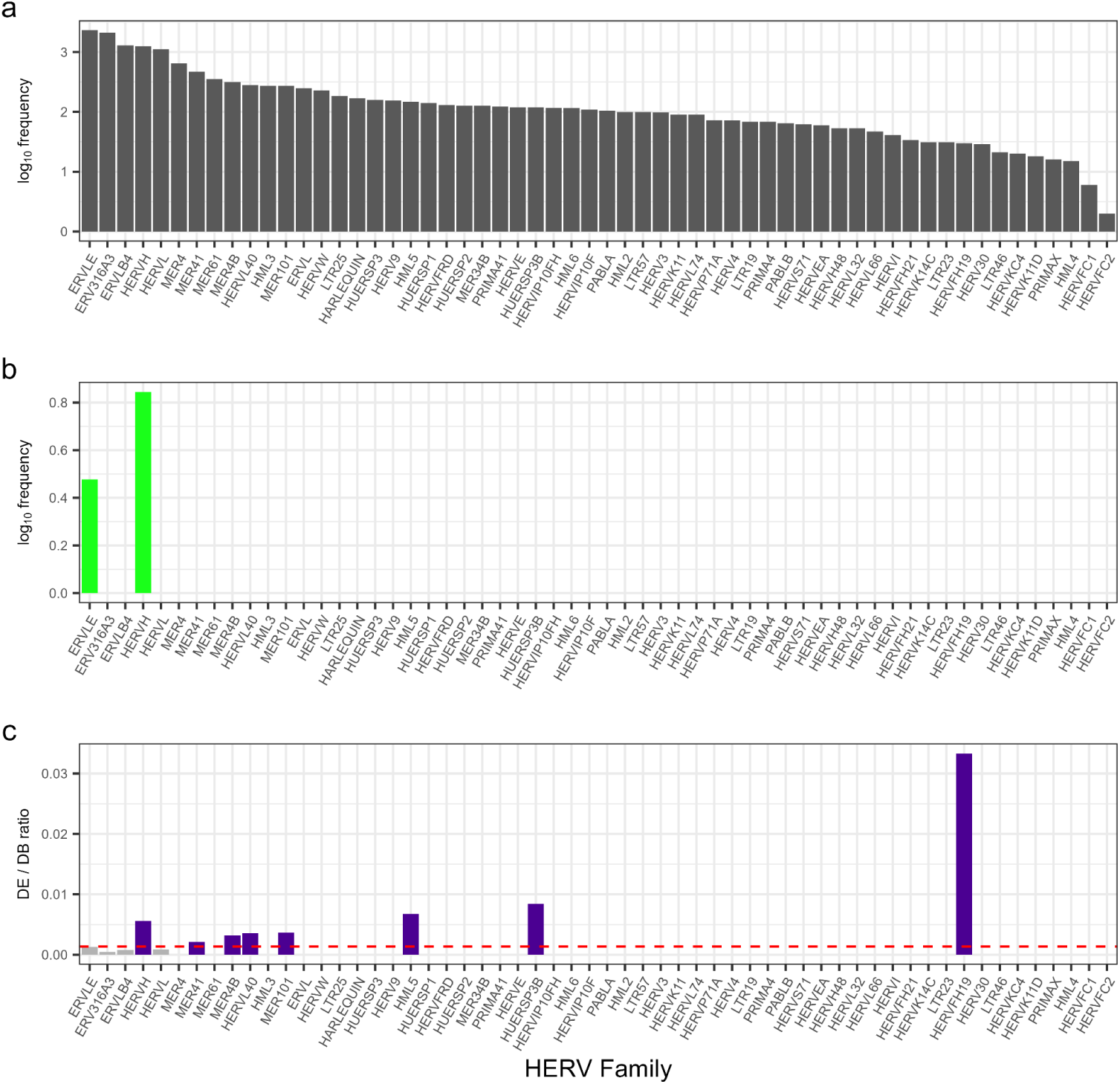
HERV frequency by family. **(a)** Log_10_ frequency of HERV families in the database (DB). **(b)** Log_10_ frequency of HERV families among the DE HERVs in our analysis (DE). **(c)** Ratio of frequency of DE HERVs in our analysis divided by frequency in the database. The dashed line indicates the expected ratio (total number of HERVs in the database divided by the total number).

### Host gene expression

Among 57,071 expressed genes, we identified 8,693 differentially expressed transcripts between PD and HC (FDR-adjusted *P* < 0.05), of which 7,086 were upregulated and 1,607 downregulated (Fig. 4; Supplementary Table 5). Initial PCA showed strong clustering driven by Y-chromosome gene expression, indicating sex was a major source of variance. After excluding Y-linked genes, samples no longer clustered by sex, and case–control groups showed substantial overlap, suggesting disease status is not the dominant contributor to global transcriptional variation (Supplementary Fig. 2). After applying apeglm shrinkage and |log_2_FC| > 0.1 cutoff, 8,027 remained significant. The most significantly upregulated genes were SPANXN1, DPP3P2, IGLV9-49, PGAM5P1, PRR35 while RNU2-36P, RNA5-8SN3, DEFA1B were among the most significantly downregulated in PD. Notably, several genes with large expression changes (|log_2_FC| > 1), such as ENSG00000281181, ENSG00000280614, and ENSG00000280800, lack HGNC annotation, suggesting the presence of potentially novel transcripts or uncharacterized non-coding RNAs.

**Fig. 4:**
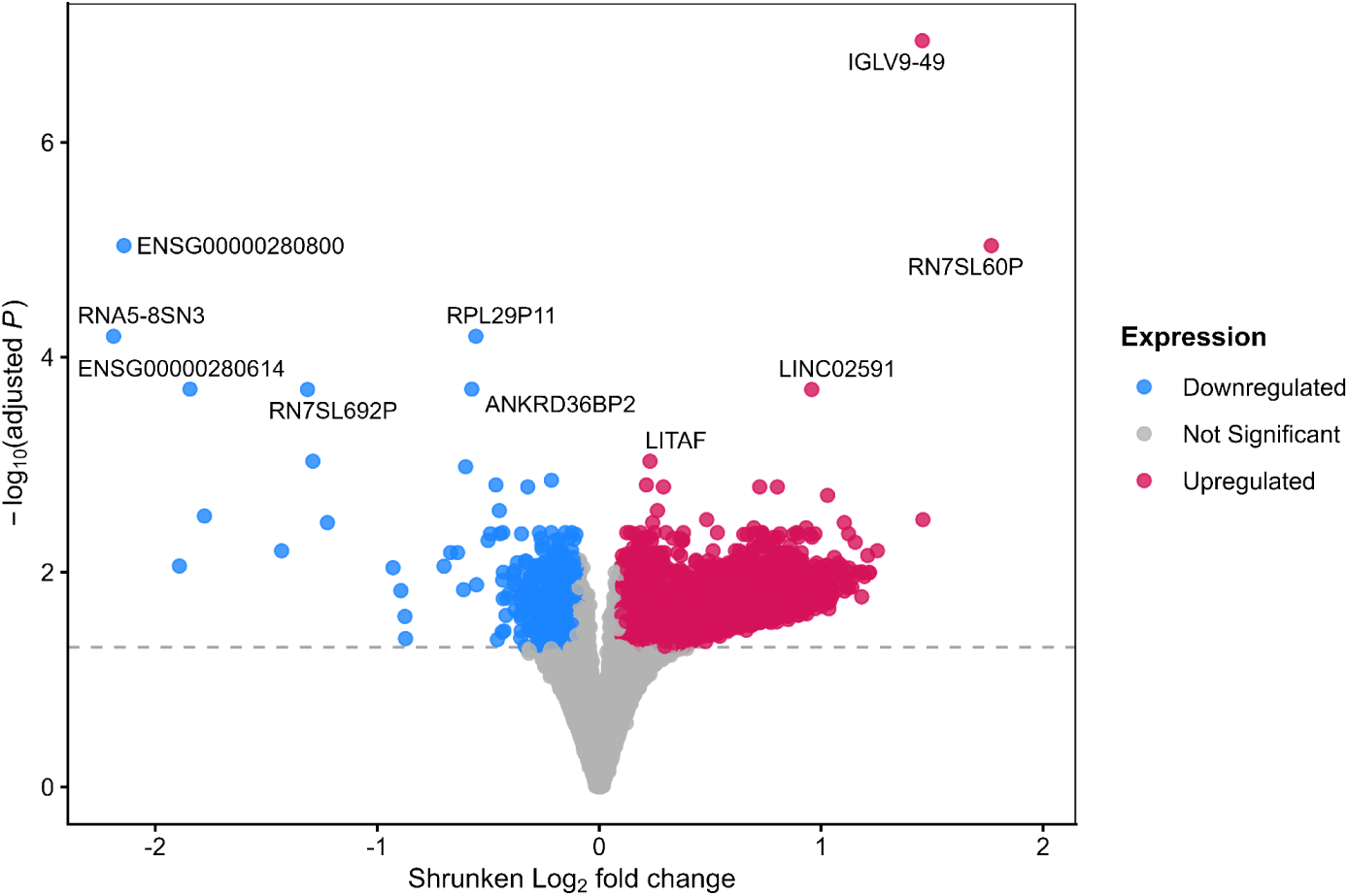
Differential gene expression analysis of all PD versus HC. Volcano plot of differential gene expression analysis in all PD compared to HC. Dots in the volcano plot represent genes, with red indicating upregulation, blue indicating downregulation and grey indicating not significant (FDR-adjusted *P* > 0.05. The top 10 most statistically significant genes ranked by adjusted *P* were selected for labeling.

### Proximal gene set enrichment and pathway analysis

Gene set enrichment analysis of HERV-proximal genes revealed multiple significantly (*P* <0.05) enriched Gene Ontology Biological Processes (GO:BP) terms in the ranked gene list (Fig. 5). 36 pathways were significantly negatively enriched (NES <0), compared to only 3 positively enriched pathways (NES > 0) at an FDR-adjusted *P* < 0.05 (Supplementary Table 6). Metabolic pathways were suppressed and reported the highest set size (n > 500) and followed by gene expression (GO:0010467), cellular response to stress (GO:0033554), RNA biosynthetic process (GO:0032774) and proteolysis involved in protein catabolic process. Cell migration (GO:0016477) and cell motility (GO:0048870) reported set sizes (n > 100) followed by actin-mediated cell contraction (GO:0070252) in activated enriched pathways. Overall, the top enriched themes (FDR-adjusted *P* < 0.05) cluster into cell movement, transcriptional reprogramming and nucleic acid metabolism.

**Fig. 5:**
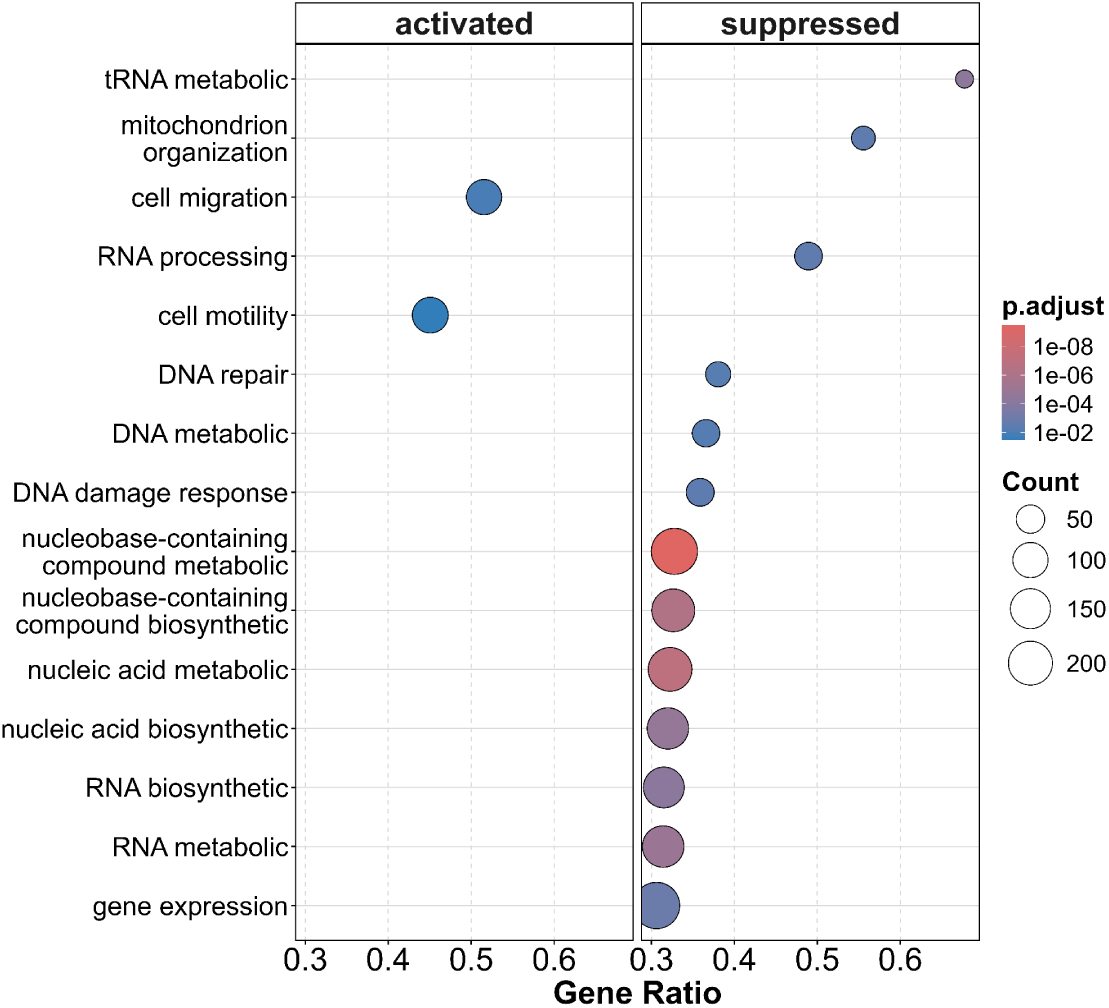
Expressed pathways of genes proximal to HERVs. The top 15 most significant pathways in all PD, ranked by adjusted *P* (p.adjust). Results are divided into two panels: activated (left) and suppressed (right). The Gene Ratio represents the proportion of genes within a given pathway that are present in the input gene list. The size of each dot reflects the absolute number of genes from the input list that are annotated to a given pathway and contribute to its enrichment score. The color of each dot encodes the statistical significance of pathway enrichment: red indicates high significance (p.adjust ≈ 1×10⁻⁸), through shades of pink and purple, to blue, indicating lower but still meaningful significance (p.adjust ≈ 1×10⁻²). The Y-axis lists the enriched Gene Ontology biological process terms identified in the analysis.

### Cell type abundance

Beta regression analysis revealed that all PD, iPD and LRRK2 carriers were significantly associated with two immune cell populations after correction for multiple comparisons (FDR < 0.05). The CIBERSORTx results are reported in Supplementary Table 7. Neutrophil proportions showed a significant positive association with all PD versus healthy controls (β ≈ 0.15), indicating that PD gene expression was associated with greater neutrophil abundance (Fig. 6).

**Fig. 6:**
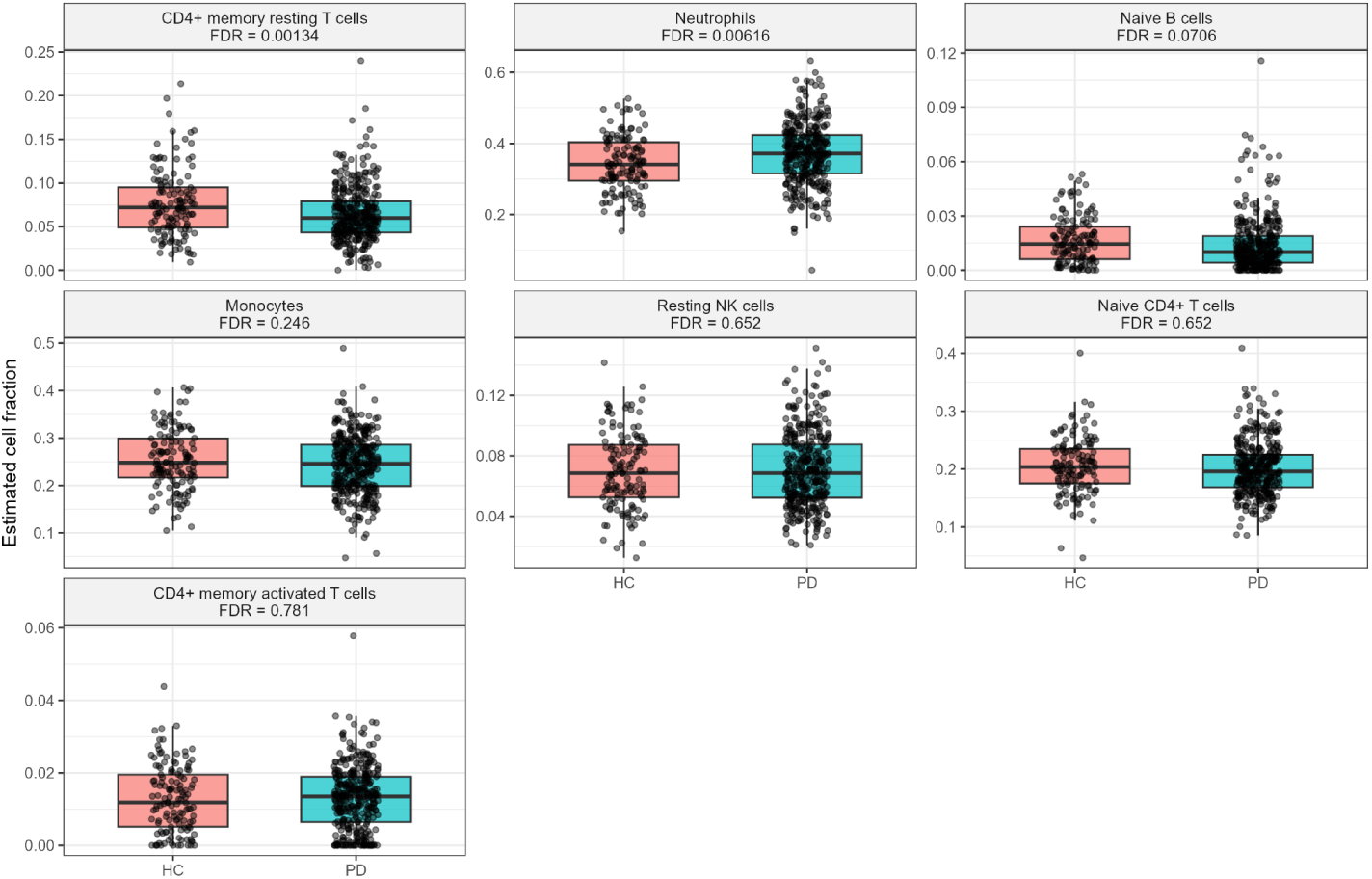
Comparison of inferred immune cell-type proportions between all PD and HC. Estimated immune cell fractions in HC (pink) and all PD (teal) by CIBERSORTx. Each dot represents an individual sample; boxplots show the median and interquartile range, with whiskers indicating variability. FDR values indicate the significance of group differences for each cell type after multiple testing corrections.

Conversely, resting CD4+ memory T cells demonstrated a significant negative association with PD (β ≈ −0.25), suggesting that PD gene expression corresponded with lower proportions of this cell type (Fig. 6). The remaining five immune cell types, resting NK cells, activated CD4+ memory T cells, naive CD4+ T cells, monocytes, and naive B cells showed no statistically significant associations with PD following FDR correction, though naive CD4+ T cells and monocytes exhibited small negative point estimates. Results were confirmed using both linear and beta regression models.

## Discussion

To our knowledge, this is the first study to investigate HERV loci in all PD and PD subtypes. We characterized the differential locus-specific expression of HERVs in PD compared to HC blood samples. 20 HERV loci were found to be significantly differentially expressed. We additionally investigated HERV loci expression patterns in iPD and LRRK2 PD compared to HC, which confirmed dysregulation in PD populations and a consistent skew towards upregulation in peripheral blood. The overlap in HERV loci suggests systematic reactivation is not only a relevant player in aging but also in disease, with HERV loci as potential biomarkers for both^36^.

The loss of HERV differential expression upon adjustment for neutrophil and resting CD4+ memory T cells proportions indicates that cell-type composition is a primary driver of the observed HERV signal in bulk blood. It is also consistent with a non-cell-autonomous model of HERV derepression, whereby the peripheral inflammatory environment drives HERV loci across blood cell populations. This pattern in neutrophils has been described in other inflammatory conditions such as systemic lupus erythematosus, where HERV-K expression in peripheral blood correlates with neutrophil activation state and interferon status rather than representing a cell-type-intrinsic transcriptional program^37^. The exception of ERVLE_8q12.3f, warrants further investigation as a candidate for cell-autonomous derepression.

HERV-H exhibits the strongest enrichment, whereas ERVLE and ERV316A3 are highly represented. Research connecting HERVs to PD has concentrated on the HERV-K family, one of the most biologically active, and one study has identified 2 loci, HERV-P and ERV316A3, in a cohort with European ancestry. Our study highlights additional candidate loci and evolutionary families, particularly the dominance of HERV-H copies, known for immunosuppressive properties in a variety of cancers and high sequence similarity leading to genomic instability^38^. HERV-H elements have also been cited for their role in neurodevelopment in autism and pathogenesis of MS, ALS, AD^2,39^. Another strength of our study is the identification of locus-specific heterogeneity within HERV families. For example, ERVLE elements exhibited mixed regulation, with one locus upregulated at 3q25 and two loci downregulated at 8q12 and 1p32. In contrast, MER elements (MER41, MER101, MER4B) were consistently downregulated across distinct q-arm loci, suggesting coordinated regional silencing. These patterns would be hidden and uninterpretable at the family level.

After effect size shrinkage and a modest fold-change threshold (|log2FC| > 0.1), most significant features remained. This reinforces the robustness of the signal, despite small effect sizes. It aligns with the expectation that PD is characterized by many modest but coordinated transcriptional perturbations. Notably, some of the most strongly differentially expressed features lack HGNC annotation. This indicates the presence of uncharacterized transcripts or non-coding RNAs. The relevance is heightened for HERVs, whose repetitive and poorly annotated nature often leads to incomplete representation in standard gene catalogs. The enrichment of such unannotated features raises the possibility that non-canonical transcripts, including HERV-derived elements, may shape the observed transcriptional landscape. Studies show that LTRs derived from HERVs are enriched for transcription factor binding sites, especially within HERV-H LTRs in pluripotency-related genes. Consistent with this, our analysis indicates possible changes in gene expression driven by LTR-mediated transcriptional regulation^10,40,41^.

Gene set enrichment analysis of genes proximal to differentially expressed HERVs reveals dysregulated pathways that play a role in PD. The pathways with the most statistically significant adjusted *P* values were linked to RNA metabolic processes, nucleic metabolic processes, DNA-templated transcription, mitochondrion organization and actin-mediated cell contraction. The pathways with the largest set size were regulation of metabolic processes and gene expression. Disruptions to actin cytoskeleton pathways have been widely reported in PD and linked to alpha-synuclein pathology, which alters actin polymerization dynamics and promotes formation of cofilin–actin aggregates. These changes impair synaptic vesicle trafficking, axonal transport, and overall neuronal function. Because actin remodeling underlies cell motility and contraction, its dysregulation further affects microglia activation and migration contributing to the recruitment of peripheral immune cells and neuroinflammatory response^42,43^. RNA metabolism and DNA-templated transcription alterations can impair mitochondrial gene expression and bioenergetic function while promoting oxidative stress and defective proteostasis. Together, these changes contribute to sustained transcriptional stress, ultimately compromising neuronal survival^44–47^.

Our immune deconvolution shows a positive association between PD and neutrophil proportions and a negative association with resting CD4+ memory T cells, consistent with a well-established pattern of peripheral immune dysregulation in PD. The elevated neutrophil signal aligns with the broader literature on the neutrophil-to-lymphocyte ratio (NLR) as an index of systemic inflammation in PD. A systematic review and meta-analysis of 20 studies confirmed significantly elevated NLR in PD patients relative to healthy controls across multiple study designs and ethnic populations^48^. Whether elevated NLR reflects a primary rise in neutrophils, a primary decline in lymphocytes, or both remains debated, and our approach suggests both arms may be independently altered at the transcriptomic level^49^. The reduction in resting CD4+ memory T cells we observed is similarly well-supported, with longitudinal DNA methylation-based immune profiling of the PPMI cohort identifying consistent decreases in resting CD4+ memory T cells in PD that deepened over time and correlated with non-motor symptom burden^46,47,45^ similarly reported reduced naive CD4+ T cells alongside increased neutrophils using CIBERSORT-based deconvolution of blood gene expression data. Taken together, our results corroborate the emerging consensus that PD peripheral immune dysregulation is characterized by innate immune activation and adaptive immune depletion.

## Conclusion

Our transcriptional analyses show that HERVs are significantly differentially expressed in all PD and PD disease subtypes when compared to HC. Our findings highlight the potential role in widespread immune dysfunction and LTR-mediated transcription regulation. Future research studies should focus on clinical translation and validation through both in vitro and in vivo experimentation for treatment or biomarker strategy based on HERV expression.

## Data availability

Raw sequencing data, alignment files and counts data for each sample are available at the LONI IDA, (https://doi.org/10.25504/FAIRsharing.r4ph5f) and Parkinson’s Progression Markers Initiative Data Repository (https://www.michaeljfox.org/news/ppmi-rna-sequencing-project).

Clinical data are available through Parkinson’s Progression Markers Initiative (https://www.ppmi-info.org/).

## Acknowledgements

PPMI – a public-private partnership – is funded by the Michael J. Fox Foundation for Parkinson’s Research and funding partners, including AbbVie, Alamar Biosciences, Aligning Science Across Parkinson’s (ASAP), Arrowhead Pharma, Arvinas, AskBio, BIAL, BioArctic, Biohaven, BlueRock Therapeutics, Bristol Myers Squibb, Calico Labs, Capsida Biotherapeutics, Critical Path Institute, DaCapo Brainscience, Denali, Edmond J. Safra Foundation, Eli Lilly, Gain Therapeutics, GE Healthcare, Genentech, GSK, Insitro, Johnson & Johnson Innovative Medicine, Lundbeck, Merck, Neumora, Neuron23, Novarti, Olink, Regeneron, Roche, Sanofi, Tenvie, UCB, Vanqua Bio, Voyager Therapeutics, The Weston Family Foundation. We are grateful to the GW High Performance Computing team for providing resources and support for this work.

## Ethics Approval and Consent

Not Applicable

## Author contributions

E.B., C.S.V., and K.A.C. conceived and designed the study. E.B. performed data generation and conducted the computational analyses. C.S.V. and K.A.C. provided guidance on data interpretation and contributed to analysis revision. E.B. drafted the manuscript. All authors contributed to result interpretation, manuscript writing, and approved the final version.

## Code availability

All custom scripts used for data analysis and pipeline are available at: https://github.com/ElizabethBanda/hervs-PD-paper

