## Supplementary Materials for "Mobile DNA Activity in Parkinson’s Disease: A Locus-Specific View of Endogenous Retroviruses"

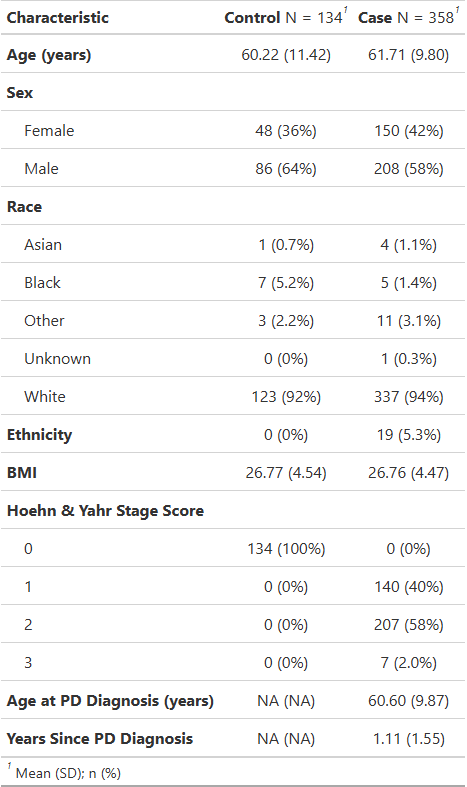

**Supplementary Table 1:** Participant Characteristics of PD and HC Cohort. This table summarizes key demographic and clinical features for the participants included in our study. Variables include age (mean + s.d.), sex, race, ethnicity, BMI, Hoehn and Yahr Stage Score, Age at PD Diagnosis and Years Since PD Diagnosis with counts and percentages also reported.

(a)
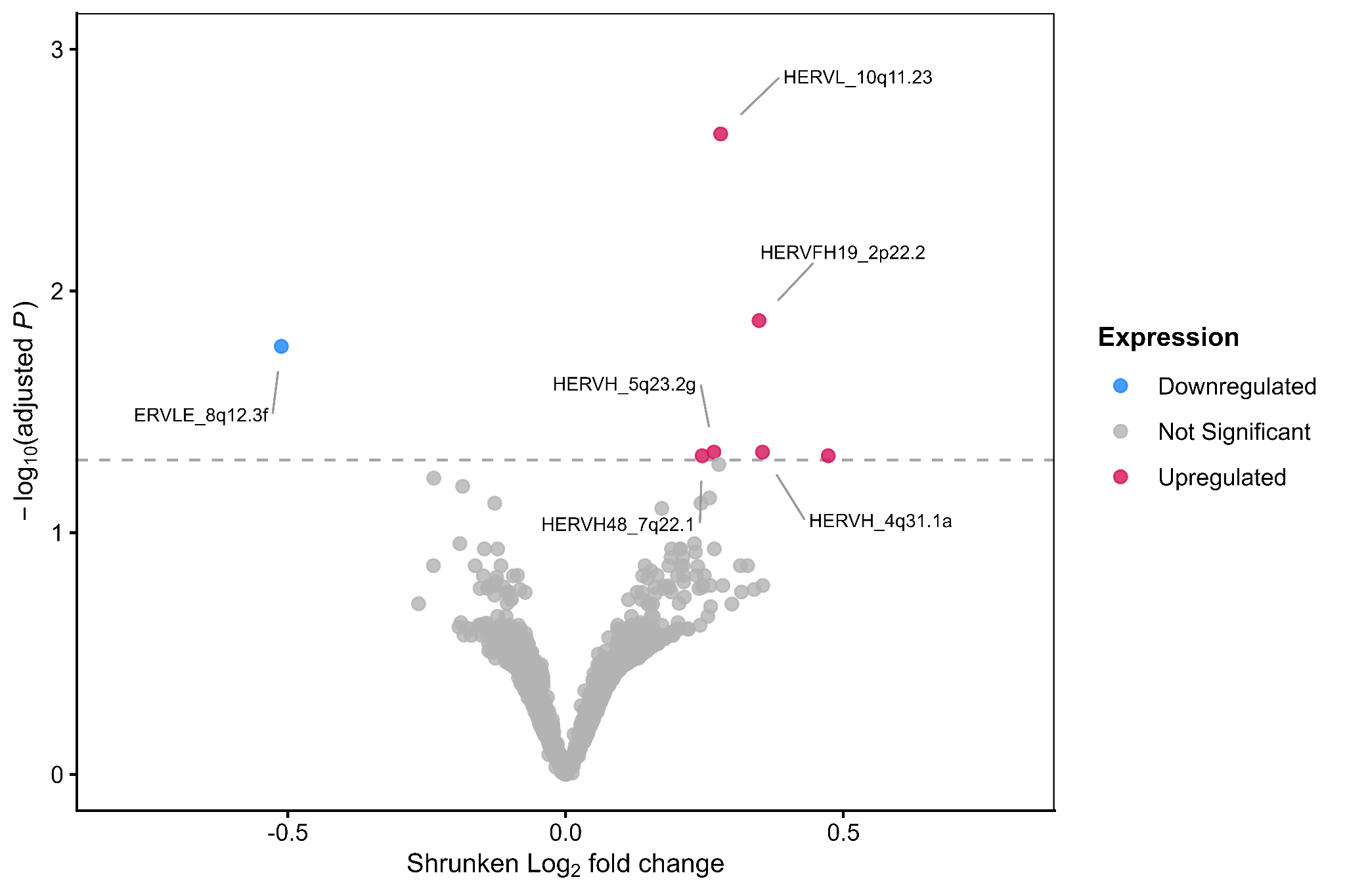

(b)
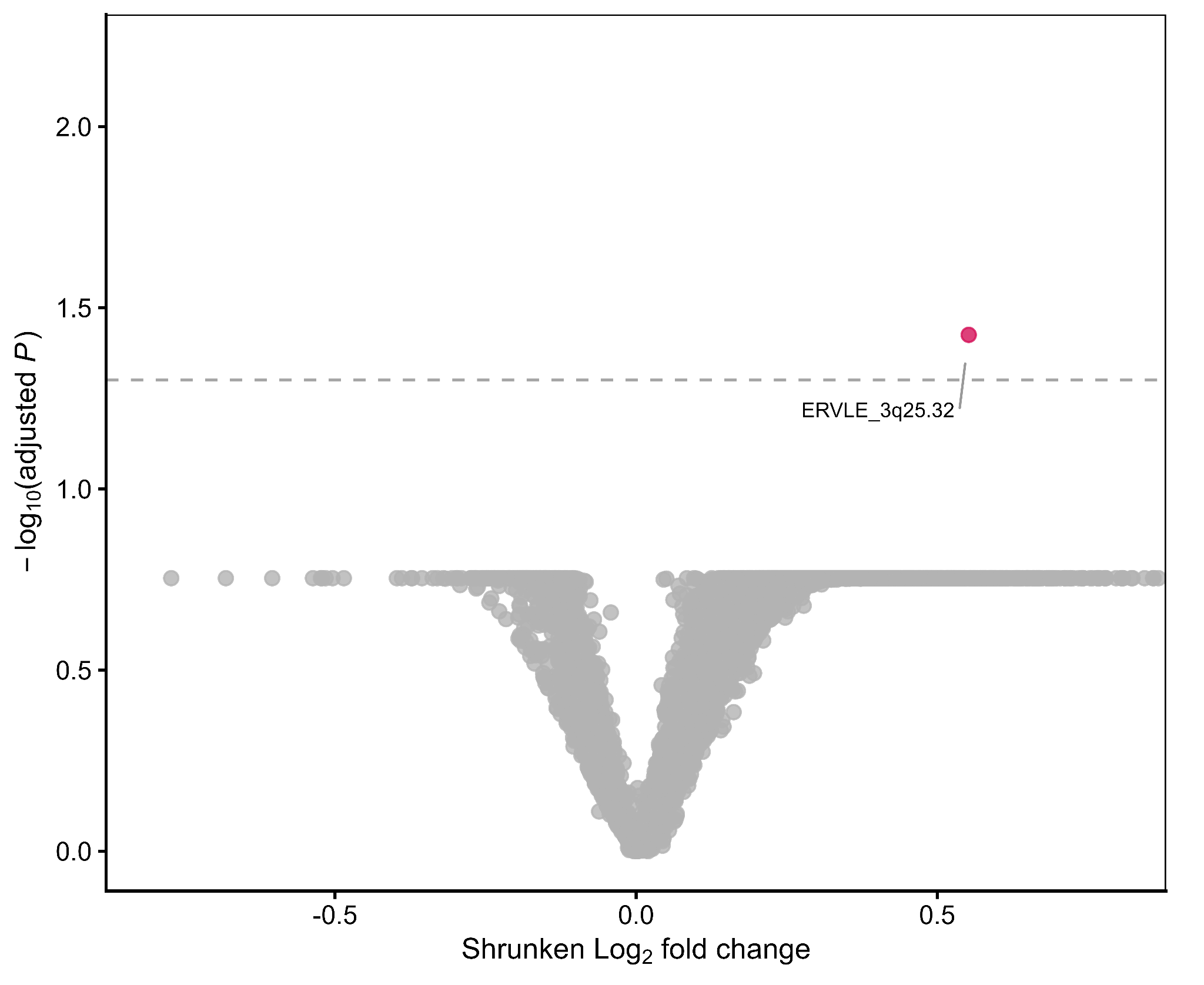

**Supplementary Figure 1 (a)** Volcano plot showing differential HERV expression in iPD compared to HC. Labeled points denote the 6 significant HERV loci in each direction (upregulated and downregulated). **(b)** Volcano plot showing differential HERV expression in LRRK2 PD compared to healthy controls. Each point represents a HERV locus; axes are as in (a). The labeled point indicates the single significantly differentially expressed locus.

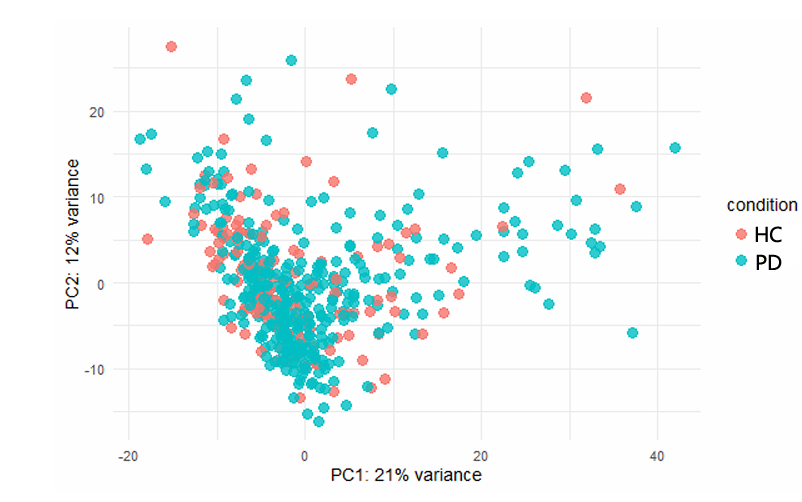

**Supplementary Figure 2:** PCA of variance-stabilized host genes using plotPCA function in DESeq2.
